# Free energy perturbations in enzyme kinetic models reveal cryptic epistasis

**DOI:** 10.1101/2025.09.05.674464

**Authors:** Karol Buda, Nobuhiko Tokuriki

## Abstract

Epistasis—the context-dependence of mutational effects—is a key driver of protein evolution, influencing adaptive pathways and functional diversity. While specific epistasis arises from direct physical interactions between mutations, non-specific epistasis emerges when a non-linear mapping links a protein’s biophysical properties to its function. Enzyme kinetic parameters are assumed to be devoid of non-specific epistasis when measured *in vitro*, enabling direct connection of epistasis to the enzyme’s structural features. Here, we show that this assumption is incorrect: enzyme catalytic parameters like *k*_cat_ and *K*_M_ inherently exhibit non-specific epistasis due to the multi-state nature of the catalytic cycle. Using *in silico* enzyme models, parameterized by free energies of ground and transition states, we simulated 1000 “mutations” or perturbations to the sub-state free energies within the kinetic ensemble. We then combined these mutations, creating one million double mutants with strictly additive free energy effects. Despite the absence of explicit mutational interactions, we observed substantial epistasis in catalytic parameters; its prevalence and complexity increasing with the number of kinetic states in the mechanism. We derived analytical conditions for the emergence of this form of epistasis in a simple kinetic model, demonstrating that non-specific epistasis depends on the relative values of key microscopic rate constants. Finally, we validated our framework by reanalyzing kinetic data for double mutants in *Bacillus cereus* β-lactamase I and found that reported specific epistasis in catalytic efficiency was substantially stronger than previously inferred, altering mechanistic interpretations. Our results identify an intrinsic, previously unknown source of epistasis that can distort both the magnitude and sign of mutational effects in enzyme kinetics. We provide theoretical and computational tools for recognizing and correcting for this form of non-specific epistasis, enabling accurate mechanistic inference from kinetic data and improving our understanding of the links between epistasis, structure-function relationships, enzyme evolution, and protein design.

**Author Summary:** Enzymes help organisms convert reactants to products through a series of steps; each associated with an energy that dictates how well the enzyme catalyzes a reaction. Enzymes evolve to become more efficient, or catalyze new reactions, through mutations that change the free energies of these steps. Sometimes, the effect of one mutation depends on the presence of another, a phenomenon called epistasis. Epistasis is typically studied by measuring the effect of mutations on standard enzyme parameters under the assumption that changes in these values reflect structural interactions between mutations in the protein. Our study shows that this assumption is misleading. Even when mutations act independently on the energies of steps in an enzyme’s reaction, when combined they can create epistasis. This phenomenon arises from the complex, non-linear relationships between the parameters that define each step in the enzyme reaction and the measurements we obtain during experiments probing enzyme functions. Using computational simulations, we mathematically derive the necessary conditions for this form of epistasis, demonstrate that epistasis increases in prevalence as the enzyme reaction becomes more complex, and apply our model to published experimental data. Our findings urge that researchers should account for these effects before drawing structural conclusions from epistasis.

## Introduction

Enzyme evolution is underpinned by the accumulation of adaptive, neo-functionalizing mutations. The effect of a mutation is dictated by the enzyme’s sequence; beneficial mutations may be contingent on specific amino acid interactions or inaccessible due to certain incompatibilities within the sequence [1–9]. This mutational context-dependence, known as epistasis, dictates the accessibility of evolutionary paths that an enzyme can take, and is a major driver of sequence diversity in nature [10]. Epistasis can dramatically change the topology of an enzyme’s fitness landscape and distort our ability to accurately assess the functional effect of a mutation across different enzymes [11–13]. Thus, our ability to predict and understand evolutionary outcomes requires a robust understanding of epistasis.

Epistasis is generally categorized using three criteria: its sign contribution relative to the effects of the individual mutations, the degree of deviation from the expected outcome relative to a wild-type (wt) reference state, and its source. When single mutational effects depart from their expected behaviour, the resulting double mutant’s function will either be greater than expected (positive epistasis) or lower than expected (negative epistasis). Upon categorizing the sign contribution of the single mutational effects, epistasis can simply amplify or diminish the mutational effects (magnitude epistasis), entirely change their sign contribution, *e*.*g*., from beneficial in one background to deleterious in another (sign epistasis), or, in the most extreme case, change the sign contribution of both interacting mutations, *i*.*e*., cause two positive mutations to elicit negative epistasis relative to the wt or *vice versa* (reciprocal sign epistasis) [14,15]. Finally, the source of epistasis is broadly categorized into two main types: specific and non-specific [16]. Specific epistasis (also called idiosyncratic epistasis [12]) arises from specific physical interactions between mutations—these may be proximal or relayed through an enzyme’s intramolecular network [17]. Functionally relevant interactions expose the intricate connections between amino acids within enzymes and help link enzyme structures to their functions [18–21]. A deep understanding of specific epistasis offers insight into the enzyme’s mechanism and provides both limitations and opportunities for changes in the enzyme’s sequence to promote existing chemical reactions or establish novel ones. However, extracting specific epistasis from functional measurements is difficult, as it is frequently distorted by non-specific epistasis.

At its core, non-specific epistasis (also referred to as global epistasis [12,22]) arises when mutational combinations appear epistatic due to some non-linear relationship between their effects on an enzyme’s biophysical property and the enzyme’s function [16,23]. The complexity of the non-specific epistasis is proportional to the complexity of the function that translates an enzyme’s biophysical parameters to the measurable, functional readout, such as enzyme catalytic activity or whole organismal fitness. The simplest example is threshold epistasis [24,25], where the relationship between protein stability and expression (which, itself, is assumed to be proportional to function) is well defined by the Boltzmann distribution—a sigmoid function relating the folded proportion of a protein and the change in free energy (Δ*G*) between folded and unfolded states. Changes in free energy, an additive trait, can result in non-additive changes to the folded fraction and thus, the observable function. Indeed, deep mutational scanning (DMS) experiments have unveiled these global sigmoidal relationships between predicted and observed functions of mutations [26–30]. Thus, a conventional practice is to estimate non-specific epistasis using sigmoidal or other similar threshold models that enact a non-linear transformation of the function to the linear free energy scale [31,32]. However, non-specific epistasis can be underpinned by more complex relationships. For example, if proteins sample more than two functionally relevant sub-states, the relationship between all sub-state Δ*G*s and the protein’s function becomes obscured.

This was conceptually elucidated by Morrison *et al*. (2021) in the form of ensemble epistasis [33]. They demonstrated that, in the presence of more than two protein sub-states, protein function can be linked to a Boltzmann weighted average of its conformational sub-states. When mutations that additively alter the free energies of each sub-state are combined, they can result in a non-additive change to the weighted average of the sub-states, resulting in complex patterns of epistasis, including sign epistasis [33]. These patterns cannot be captured by a threshold model, as the protein’s phenotype is an aggregate of several non-linear functions.

Enzyme kinetics may be another example where non-specific epistasis can arise from the ensemble of sub-states. The enzyme catalytic cycle must go through several energetic states, *e*.*g*., the enzyme-substrate complex and the enzyme-transition state (TS) complex. The relationship between the energetic level of each state and the commonly measured kinetic parameters (*k*_cat_, *K*_M_, and catalytic efficiency) can be multivariate and scales in complexity with the enzymatic mechanism. Moreover, mutations can differentially affect the relative free energies of these sub-states, and thus, multiple mutational effects can further obscure the relationship between the sub-state energies and the measured kinetic parameters. In this study, we aimed to uncover an untapped source of non-specific epistasis in enzymes. By assigning free energies to ground and transition states within an *in silico* catalytic cycle, and simulating mutations that perturb these free energies, we investigated whether additive free-energy changes between mutational combinations may create non-specific epistasis and provide a theoretical rationale for our observations. We also explored how increasing complexity in the catalytic cycle results in more non-specific epistasis. Finally, we applied our model to experimental data and extracted non-specific epistasis in the catalytic cycle from a β-lactamase double mutant. Our work demonstrates that catalytic parameters appear to be inherently epistatic and cautions researchers from making direct structural claims using epistasis obtained from enzyme kinetics measurements.

## Results

### Simple kinetic model reveals emergent epistasis

To determine whether mutations with additive effects on energetic terms may create non-specific epistasis, we established a model with no explicit interactions where a reaction coordinate is based on a hypothetical enzyme reaction governed by the classical Michaelis-Menten enzyme mechanism (**Fig. 1a**) [34]. The mechanism is defined by the following rate constants: a reversible binding step defined by rate constants *k*_1_ and *k*_-1_, as well as an irreversible chemical step of product formation, *k*_2_, hereafter referred to as “microscopic rate constants.” To obtain the microscopic rate constants of this hypothetical enzyme reaction, we assigned relative Gibbs free energies (*G*), in kcal mol^-1^, to each state in the reaction coordinate: the ground states, *i*.*e*., the enzyme and substrate (E + S) and the enzyme-substrate complex (ES), as well as the transition states, *i*.*e*., the binding transition state (E + S)^‡^ and the chemical transition state (ES)^‡^ **(Fig. 1a)**. We then calculated each rate constant using the Arrhenius equation, employing transition state theory for the approximation of each pre-exponential factor (see **Methods**). Finally, we calculated the kinetic parameters: the substrate binding dissociation constant (*K*_D_), turnover number (*k*_cat_), the Michaelis-Menten constant (*K*_M_), and catalytic efficiency (*k*_cat_ / *K*_M_), as defined by the enzyme mechanism using the simulated microscopic rate constants. The values for relative free energies (**Fig. 1a**) were chosen arbitrarily, with the requirement that the extrapolated *k*_cat_, *K*_M_, and *k*_cat_ / *K*_M_ from the computed microscopic rate constants (**Table 1**) were within the range of experimentally measured, median kinetic parameters for natural enzymes outlined in studies by Bar-Even *et al*. (2011) [35] and Copley *et al*. (2022) [36].

**Table 1.** Rate constants selected for the WT *in silico* state.

| $k_{on} (M^{-1} s^{-1})$ | $k_{off} (s^{-1})$ | $K_D (\mu M)$ | $k_{cat} (s^{-1})$ | $K_M (\mu M)$ | $k_{cat}/K_M (M^{-1} s^{-1})$ |
| --- | --- | --- | --- | --- | --- |
| $2.88 \times 10^5$ | 62.1 | 215 | 11.5 | 255 | $4.50 \times 10^4$ |

**Figure 1.**
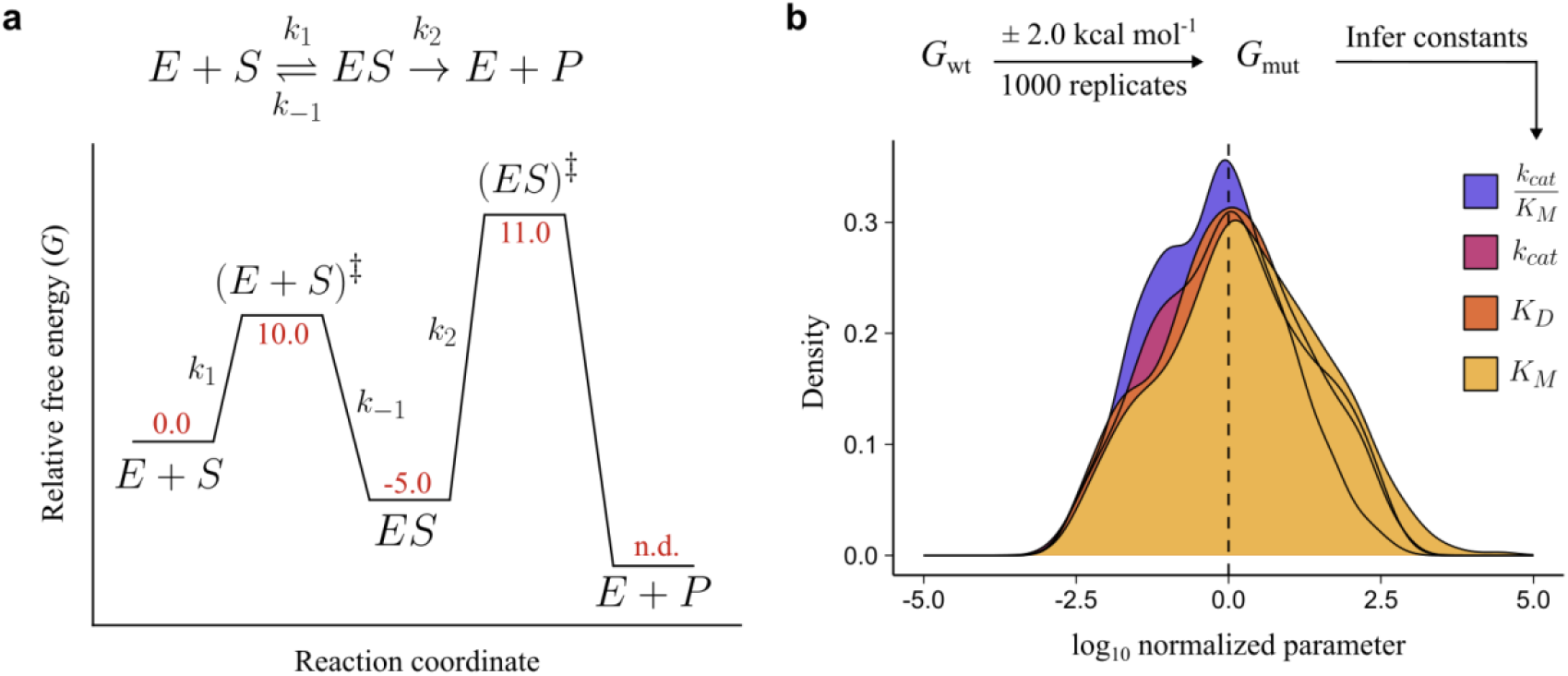
Inferred enzyme constants across 1000 *in silico* mutations are log-normally distributed. **a**, Overview of the mechanism and reaction coordinate of each state in the enzyme kinetic cycle. Red values represent Gibbs free energy for each state. E + P free energy was not defined (n.d.) as it is not required in rate constant calculation (see **Methods**). **b**, Distribution of inferred constants after log-transformation and normalization to the wt value.

Next, we simulated 1000 in silico “mutations” which are denoted by a perturbation to the free energies of each state within the reaction coordinate. A mutation perturbed each of the five sub-state’s *G* in the ensemble, both ground and transition states, by a random value between –2 and 2 kcal mol^-1^, sampled from a uniform distribution (**Fig. 1a**). The new sub-state *Gs* were recorded for each of the 1000 mutants and used to calculate new rate constants, and kinetic parameters, for the mutants. As expected, we found that the distributions of the mutants’ kinetic parameters were log-normally distributed (**Fig. 1b**). We then extrapolated a set of double mutants by summing the wt *G* to the two mutants’ Δ*G* values for each state, resulting in 10^6^ (1000 x 1000) double mutants; rate constants and kinetic parameters were calculated for each mutant. We also computed the double mutants’ predicted values for each individual rate constant and kinetic parameter by multiplying the wt value with the corresponding fold-changes in each constituent single mutant–a commonly employed null model that assumes additive mutational effects (see **Methods**). All mutant data are made available in a public repository [37].

As we did not explicitly introduce mutational interactions into our model, we aimed to probe whether differences in the double mutants’ predicted rate constants *versus* observed rate constants revealed non-specific epistasis. To account for the presence of noise in experimentally obtained rate constant measurements, we applied a 1.5-fold significance threshold for epistasis detection, which we have used previously [11]. We found that of the 10^6^ *in silico* variants, 39.4% showed significant epistasis in *k*_cat_ / *K*_M_ (**Fig. 2a**), with 28.6% at 2-fold, 7.1% at 5-fold, and 2.1% at 10-fold significance thresholds. For the 1.5-fold threshold, the majority of the epistasis was magnitude epistasis (33.0%), with some mutants exhibiting sign (5.6%) and reciprocal sign (0.8%) epistasis (**Fig. 2a**). Among all observed epistasis, both positive and negative epistasis appeared: 59.9% (117,680/196,355) positive versus 40.1% (78,675/196,355) negative. We observed exactly inverse positive-negative ratios for epistasis in the *K*_M_ (**S1 File**), but similar magnitude and sign distribution (**Fig. 2b and S1 File**). However, we did not see any epistasis in *k*_cat_ (**Fig. 2c**) or *K*_D_ (**Fig. 2d**). The statistics for all kinetic parameters can be found in **S1 File**.

**Figure 2.**
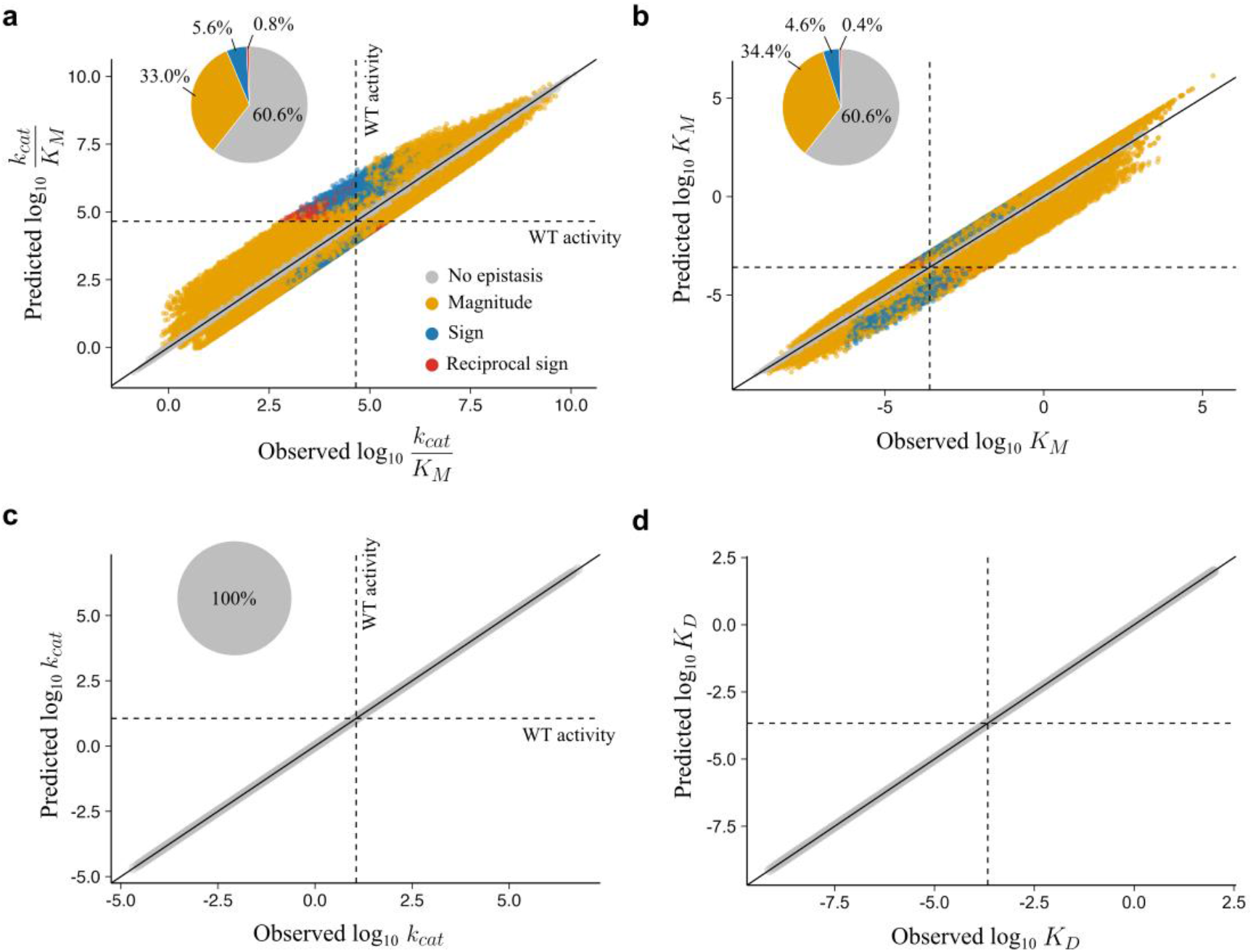
Non-specific epistasis in catalytic efficiency arises from the null model. Correlation of observed and predicted mutational effects with listed proportions of magnitude, sign, and reciprocal sign epistasis for the **a**, catalytic efficiency (*k*_cat_ / *K*_M_) **b**, turnover number (*k*_cat_), **c**, Michaelis-Menten constant (*K*_M_) and **d**, enzyme-substrate dissociation constant *K*_D_.

### Mechanisms for non-specific epistasis in a simple kinetic ensemble

The emergent epistasis from free energy perturbations in enzyme kinetics cannot be easily captured by a non-linear transformation (**Fig. 2a**) such as a sigmoidal model or other threshold model, as is often the case for non-specific epistasis. These observations suggest that mutational behaviors in kinetic parameters are more complex compared to other biophysical properties. Thus, we conducted a theoretical exploration by dissecting how epistasis arises in *K*_M_ (**Fig. 2b**). We define epistasis in *K*_M_ as deviation from a multiplicative null model, thus:

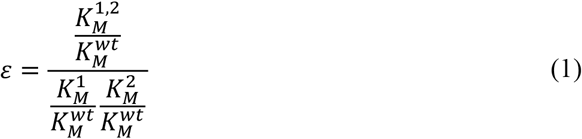

And since *K*_M_ is defined by the microscopic rate constants *k*_-1_, *k*_1_, and *k*_2_, a single mutational effect on *K*_M_ is simply the fold-change that a mutation elicits on each of the rate constants:

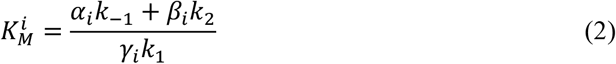

Where *i* is the relative mutational number (mutation one or mutation two), *α* is the mutation’s fold-change on *k*_-1_, *β* is the mutation’s fold-change on *k*_2_, and *γ* is the mutation’s fold-change on *k*_1_. Thus, the double mutational effect is:

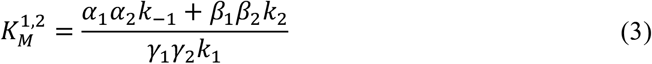

Upon substituting **eq. 1** with **eq. 2, 3, and 11 (see Methods)** and setting epistasis to zero (see **S2 File** for details), we found that:

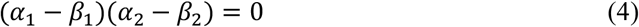

In other words, non-specific epistasis in the simple kinetic ensemble only arises when each of the two mutations has a differential effect on *k*_-1_ (*α*) and effect on *k*_2_ (*β*). Using a similar approach, we expand on the theoretical rationale for the absence of epistasis in *k*_cat_ and *K*_D_ in **S2 File**.

Using our starting wt rate constants (**Table 1**), we computed the expected magnitude of epistasis in *K*_M_ for mutational effects on *k*_-1_ (*i*.*e*., *α*_1_ for mutation one and *α*_2_ for mutation two) and *k*_2_ (*i*.*e*., *β*_1_ for mutation one and *β*_2_ for mutation two). We found that negative epistasis arose when mutations had opposite effects to each other, *e*.*g*., one mutation increased *α*_i_ relative to *β*_i_ while the other mutation decreased the ratio or *vice versa* (quadrants B and D in **Fig. 3a**). Positive epistasis, on the other hand, arose only when both mutations decreased *α*_i_ relative to *β*_i_ (quadrant A in **Fig. 3a**), but not when both mutations increased the ratio, in which case no significant epistasis was observed (quadrant C in **Fig. 3a**). We provide examples of how mutations can perturb sub-state free energies to produce double mutants that lie across different quadrants (**Fig. 3b**). We also found that our simulated dataset of randomly sampled mutational combinations fit these observations, as expected, and variants with magnitude, sign, and reciprocal sign epistasis were found in quadrants A, B, and D in **Fig. 3c** (with quadrant C populated with variants that show no significant epistasis; **Fig. 3c**). Moreover, we found that the relationship between the *α*_i_ / *β*_i_ and positive epistasis is sensitive to the starting value of *k*_-1_ (**Fig. 3d**). When *k*_-1_ < *k*_2_, positive epistasis was only observed when both mutations increased the *α*_i_ / *β*_i_. When *k*_-1_ = *k*_2,_ no positive epistasis was observed. When *k*_-1_ > *k*_2_ (equivalent to our simulated conditions), positive epistasis was only observed when both mutations decreased the *α*_i_ / *β*_i_. Furthermore, the strength of positive epistasis increased as *k*_-1_ became much greater than or much less than *k*_2_ (**Fig. 3d**). Thus, the relative values of the key rate constants, as well as the degree to which mutations modulate them, are sufficient parameters for prediction and understanding of non-specific epistasis in kinetic parameters for simple enzyme mechanisms.

**Figure 3.**
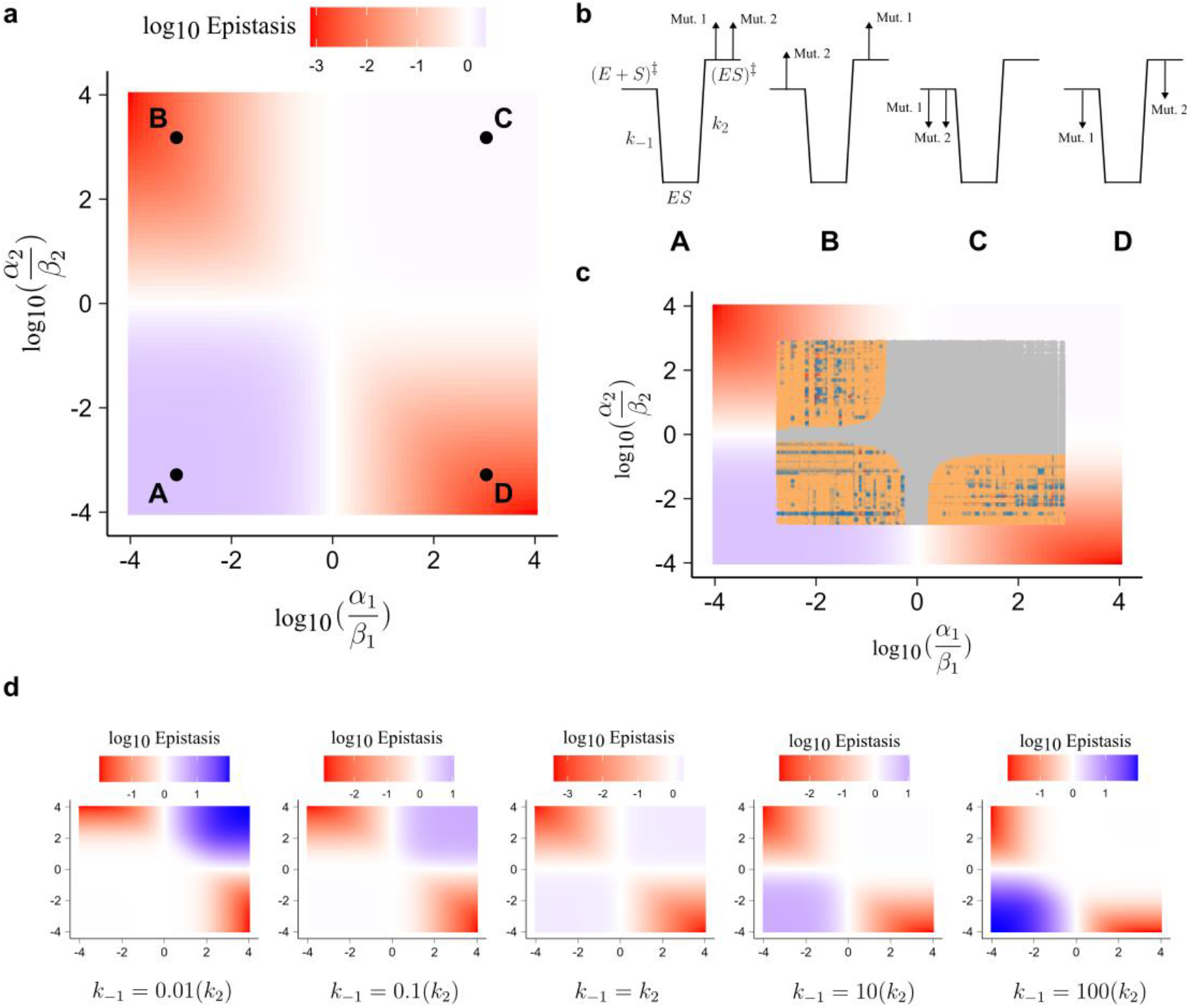
Mutational effects on *k*_-1_ and *k*_2_ are sufficient to account for non-specific epistasis in the kinetic ensemble. **a**, The observed non-specific epistasis as a function of ratios of mutations 1 effect on *k*_-1_ (*α*_1_) and *k*_2_ (*β*_1_) versus mutations 2 effect on *k*_-1_ (*α*_2_) and *k*_2_ (*β*_2_). Four extreme examples (A–D) are shown and explored in **b. b**, Four non-exhaustive examples of mutational effects that could lead to the observed patterns of epistasis in **a. c**, Distribution of simulated data from **Fig. 2b** overlayed on the relationship shown in **a. d**, The change in observed non-specific epistasis with variation in starting values of *k*_-1_ in the wt enzyme.

### Increasing kinetic model complexity amplifies epistasis

If epistasis can arise in a simple kinetic model and appears in kinetic parameters that are defined by the sum of multiple microscopic rate constants (**S2 File**), how does increasing the reaction mechanism complexity influence non-specific epistasis? To address this question, we established a reaction coordinate based on a more complex mechanism with a reversible chemical step, as well as an irreversible product release step (**Fig. 4a**). The established reaction coordinate was rate-limited by the chemical step. We retained relative energies for states from the previous model and introduced energies for the new states. We note that these changes lowered the values for *k*_cat_ and *K*_M_ such that they deviate from median reported values across enzymes [35,36], although *k*_cat_ / *K*_M_ was almost equivalent (**Table 2**). As with the previous model, we simulated 1000 mutations and monitored the emergent kinetic parameter distributions–these were also log-normally distributed (**Fig. 4b**), albeit we found that the shape of the distributions, particularly that of the *K*_M_, was broader than the simple model, likely owing to more complex equations underlying the kinetic parameters (see **Methods**).

**Table 2.** Rate constants selected for the WT *in silico* state in the complex model.

| $k_{\text{on}} (\text{M}^{-1} \text{s}^{-1})$ | $k_{\text{off}} (\text{s}^{-1})$ | $K_D (\mu\text{M})$ | $k_{\text{cat}} (\text{s}^{-1})$ | $K_M (\mu\text{M})$ | $k_{\text{cat}}/K_M (\text{M}^{-1} \text{s}^{-1})$ |
| --- | --- | --- | --- | --- | --- |
| $2.88 \times 10^5$ | 62.1 | 215 | 0.38 | 8.6 | $4.37 \times 10^4$ |

**Figure 4.**
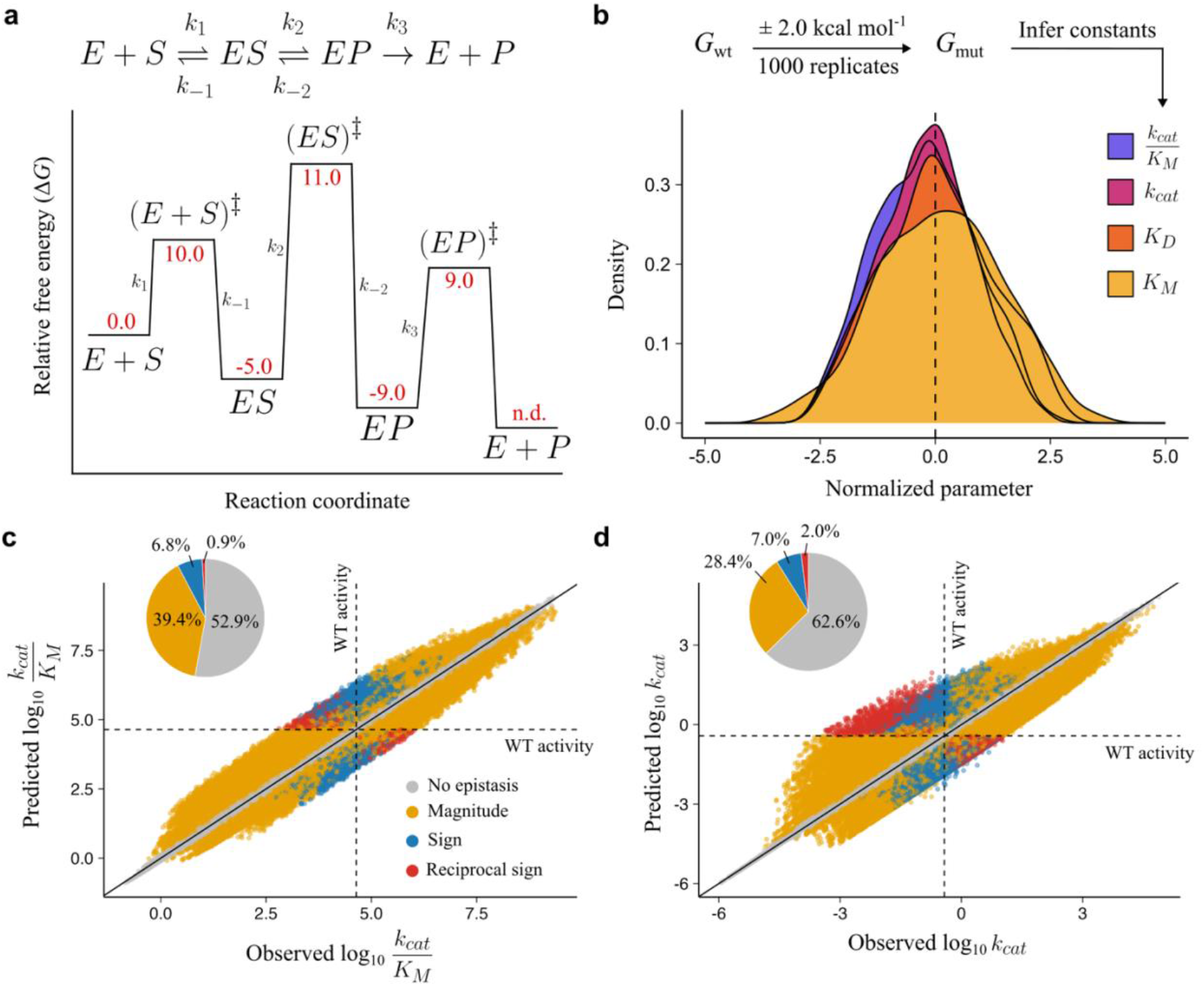
Non-specific epistasis in catalytic efficiency increases with a more complex mechanism. **a**, Overview of the mechanism and reaction coordinate of each state in the enzyme kinetic cycle. Red values represent Gibbs free energy for each state. E + P free energy was not defined (n.d.) as it is not required in rate constant calculation (see **Methods**). **b**, Distribution of inferred constants after log-transformation and normalization to the wt value. **c, d**, Correlation of observed and predicted mutational effects with listed proportions of magnitude, sign, and reciprocal sign epistasis for **c**, catalytic efficiency (*k*_cat_ / *K*_M_) and **d**, *k*_cat_.

As previously, *K*_D_ did not show non-specific epistasis (**S3 File**). In addition to epistasis observed for *K*_M_ (**S3 File**) and *k*_cat_ / *K*_M_, we also observed non-specific epistasis in *k*_cat_. The proportion of epistasis in *k*_cat_ / *K*_M_ grew from 39.4% in the simple model to 47.1% (**Fig. 4c**). The increase in epistasis was found across all forms: magnitude epistasis (39.4%), sign (6.8%) and reciprocal sign (0.9%) epistasis (**Fig. 4c**). Like in the simple model, most of the significant epistasis was positive: 60.9% (143,060/234,976) positive versus 39.1% (91,916/234,976) negative. The epistasis in *k*_cat_ was lower (37.4%), though the proportion of sign (7.0%) and reciprocal sign (2.0%) was greater than that of *k*_cat_ / *K*_M_ in both models (**Fig. 4d**). Furthermore, *k*_cat_ epistasis appeared less skewed to positive effects than *k*_cat_ / *K*_M_: 51.5% (95,842/186,050) positive vs 48.5% (90,208/186,050) negative. The statistics for all kinetic parameters can be found in the **S3 File**. All mutant data are made available in a public repository [37].

As expected, and outlined in the **S2 File**, unlike the simple model, we observed epistasis in *k*_cat_ because it included a summation in its functional mapping to its constituent microscopic rate constants (**eq. 4.1** in **Methods**). The equation for *K*_D_, however, does not change with increasing complexity of the enzyme mechanism and therefore remains non-epistatic.

### Identification of epistasis from the catalytic cycle of Bacillus cereus β-lactamase I

Finally, we aimed to demonstrate that epistasis within the catalytic cycle is found in experimental data. Thus, we sought out to find literature that met the following criteria: (*i*) an explicit statement of the enzyme mechanism, (*ii*) the acquisition of all microscopic rate constants that are constituents of *k*_cat_ and *K*_M_, and (*iii*) the availability of all measurements for the wt, both single mutants, and the double mutant. We found one study which met our requirements–an investigation into K73R and E166D in *B. cereus* β-lactamase I by Gibson and Waley (1990) [38]. We note that two values (*k*_3_ for K73R and *k*_3_ for K73R/E166D) were reported as thresholds rather than discrete measurements; we simply used the value provided as the threshold for convenience.

To extract epistasis, we compared the predicted values of *k*_cat,_ *K*_M_, and catalytic efficiency obtained from the fold-changes in the microscopic rate constants *versus* the measured kinetic parameters. First, to ensure microscopic rate constants could be accurately used to compute the kinetic parameters, we determined whether the calculated values for the kinetic parameters in each variant matched the measured values. We found that six out of twelve computed parameters showed greater than 1.5-fold error to the measured values, suggesting that kinetic parameter computation from microscopic rate constants already introduces a substantial degree of error that may distort predictions (**Table 3**). Thus, to ensure that the error was systematically represented in calculations of epistasis, we ignored the experimentally measured kinetic parameters and exclusively used the computed kinetic parameters to obtain fold-changes for each mutant (see **Methods**).

**Table 3.**
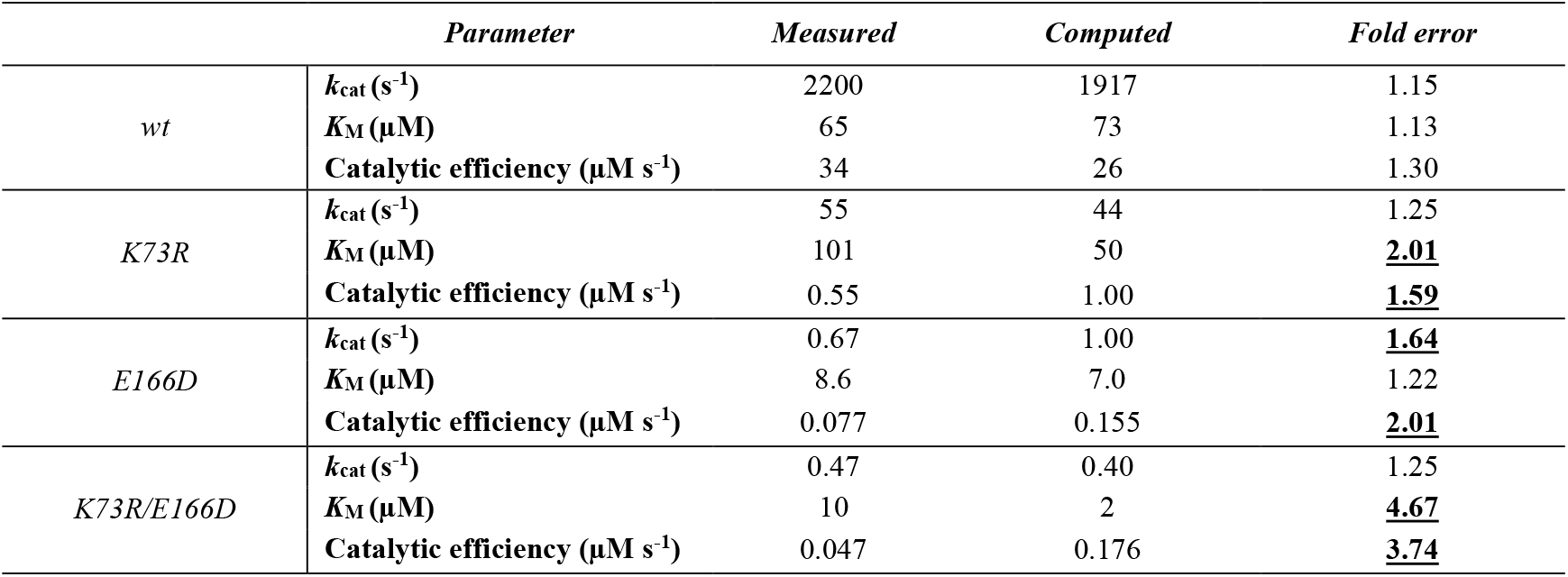
Errors associated with computation of kinetic parameters from rate constants *versus* the measured kinetic parameters Gibson and Waley (1990)

|  | *Parameter* | *Measured* | *Computed* | *Fold error* |
| --- | --- | --- | --- | --- |
| *wt* | $k_{cat}$ (s<sup>-1</sup>) | 2200 | 1917 | 1.15 |
| | $K_M$ (μM) | 65 | 73 | 1.13 |
|  | Catalytic efficiency (μM s<sup>-1</sup>) | 34 | 26 | 1.30 |
| *K73R* | $k_{cat}$ (s<sup>-1</sup>) | 55 | 44 | 1.25 |
| | $K_M$ (μM) | 101 | 50 | **<u>2.01</u>** |
|  | Catalytic efficiency (μM s<sup>-1</sup>) | 0.55 | 1.00 | **<u>1.59</u>** |
| *E166D* | $k_{cat}$ (s<sup>-1</sup>) | 0.67 | 1.00 | **<u>1.64</u>** |
| | $K_M$ (μM) | 8.6 | 7.0 | 1.22 |
|  | Catalytic efficiency (μM s<sup>-1</sup>) | 0.077 | 0.155 | **<u>2.01</u>** |
| *K73R/E166D* | $k_{cat}$ (s<sup>-1</sup>) | 0.47 | 0.40 | 1.25 |
| | $K_M$ (μM) | 10 | 2 | **<u>4.67</u>** |
|  | Catalytic efficiency (μM s<sup>-1</sup>) | 0.047 | 0.176 | **<u>3.74</u>** |

First, we found significant, specific epistasis between K73R and E166D in *B. cereus β*-lactamase I. In all cases, the double mutant exhibited diminishing losses in the microscopic rate constants due to positive epistasis: 21.0-fold for *k*_1_, 6.3-fold for *k*_-1_, 16.7-fold for *k*_2_, and 13.5-fold for *k*_3_. Next, we found weak, but significant, non-specific epistasis in *K*_M_ and catalytic efficiency, but none in *k*_cat_ (**Fig. 5a-c**). This means that even in the absence of specific epistasis seen in the microscopic rate constants, we would have observed non-specific epistasis leading to 2.2-fold greater *K*_M_ and a 2.4-fold lower catalytic efficiency than predicted. In fact, correcting for non-specific epistasis reveals a stronger contribution of specific epistasis than calculated from the null model: the apparently neutral 1.3-fold epistasis in *K*_M_ is corrected to 3.0-fold, and the 37.7-fold positive interaction between K73R and E166D in catalytic efficiency results in a corrected 90.6-fold improvement. Although the contribution of non-specific epistasis for these mutations is relatively weak to the effect of specific epistasis, its correction allows for a superior approximation of the greater contribution to diminishing losses in function by the K73R/E166D mutational interaction. Thus, our method is able to extract specific epistasis and, in the case of K73R/E166D *B. cereus* β-lactamase I, it is 2.4-fold stronger in diminishing losses of catalytic efficiency than previously thought.

**Figure 5.**
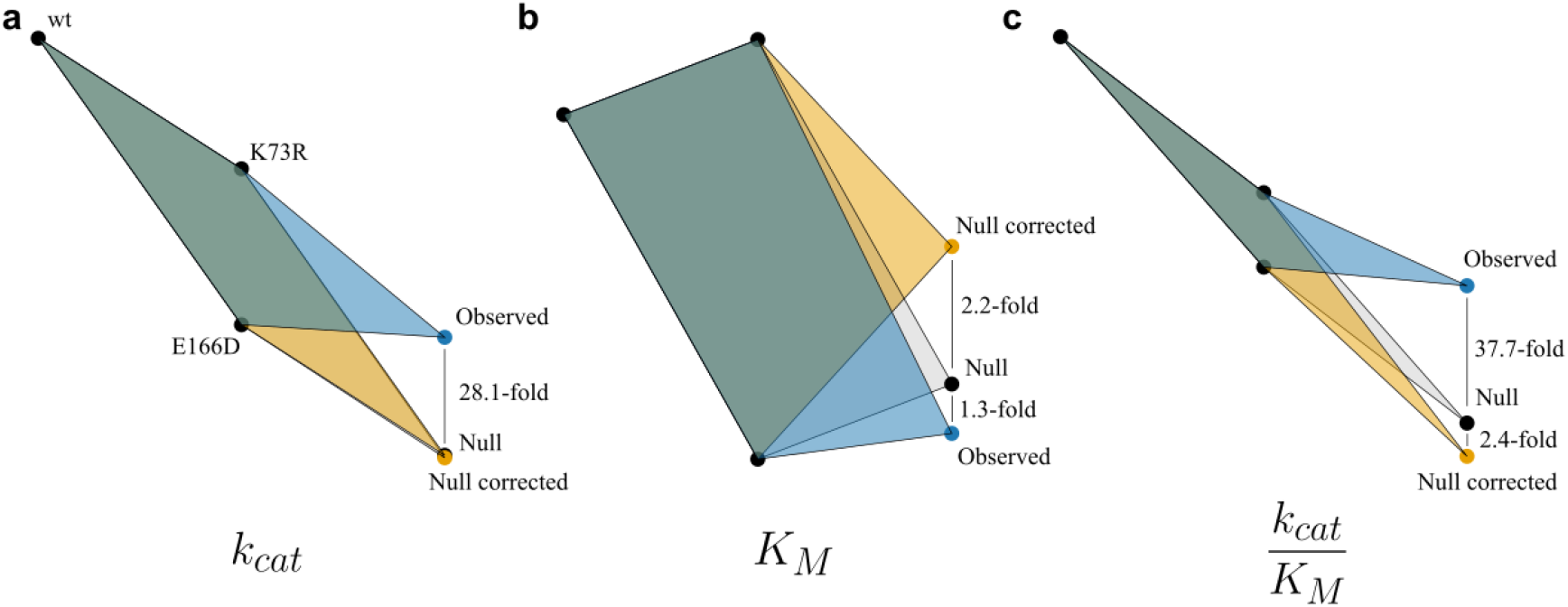
Correcting for non-specific epistasis in the null model exposes stronger positive effects. The observed and predicted fold-changes in kinetic parameters of the double mutant based on single mutational effects using a null model and one corrected from non-specific epistasis in the catalytic cycle for **a**, computed *k*_cat_. **b**, computed *K*_M_ and **c**, catalytic efficiency of K73R/E166D in *B. cereus* β-lactamase I.

## Discussion

In this study, we uncovered a novel form of epistasis, intrinsic to the catalytic cycle, that can distort predictions of a mutant’s kinetic parameters. By comparing two *in silico* models of an enzyme reaction, we demonstrated how increasing complexity in the enzyme mechanism can result in more non-specific epistasis in the double mutants. This form of epistasis can be successfully identified in experimental data, albeit it requires deep functional annotation of the enzyme’s microscopic rate constants.

Our findings necessitate a more critical evaluation of epistasis calculated from enzyme kinetic parameters. Measurements obtained from assays conducted on a purified enzyme are often assumed to be exempt from the influence of non-specific epistasis, as the confounding mutational effects on the physical properties of the enzyme can be controlled through appropriate experimental design. Likewise, the existence of a complex ensemble is often ruled out if enzyme progress curves do not exhibit notable burst or lag kinetics [39]; thus, one may assume that ensemble epistasis is also absent. Hence, many studies have used fold-changes in kinetic parameters of mutants–catalytic efficiency in particular–to connect the changes in structural features of the mutants to the emergent, specific epistasis [11,19,40–42]. Here, we have shown that non-specific epistasis in the catalytic cycle has the potential to distort the contribution of specific epistasis, not just in magnitude but also in sign. Indeed, it is very unlikely that kinetic parameters such as catalytic efficiency and the Michaelis-Menten constant will be free of non-epistasis, as that would at the very least require identical modulations of the binding and catalytic transition states in the enzyme mechanism. This form of non-specific epistasis is likely to be accentuated by changes in the rate-limiting steps for the inferred kinetic parameter during evolution, and shifts in these values will reveal strong epistasis–something we have observed experimentally [40]. Finally, it is in fact rare for enzymes to proceed through a simple Michaelis-Menten mechanism, as many enzymes rely on intermediates or multiple products and/or cofactors, further contributing the non-specific epistasis that arises. Thus, we expect that non-specific epistasis in kinetic ensembles is pervasive.

A deep exploration of non-specific epistasis, particularly its prevalence, patterns, and strength across enzymes, requires a rigorous investigation of each enzyme’s mechanism and the characterisation of all microscopic rate constants for the single and double mutants of interest. The stringency of these requirements likely accounts for the scarcity of studies that meet the necessary criteria, as well as the general unawareness of the existence of non-specific epistasis in the catalytic cycle. Collecting the necessary data to explore this form of epistasis is experimentally challenging, requiring expertise in enzymological techniques [43–45]. However, it is a necessary step in formulating accurate hypotheses regarding the impact that non-specific epistasis plays in enzyme evolution and design. Furthermore, it is essential for efforts aiming to directly connect structural features within enzyme mutants to the emergent epistasis that arises in functional measurements, as non-specific epistasis obscures this relationship. We therefore encourage researchers to either abstain from implicating structural features of mutants in their explanations of the sources of epistasis in kinetic parameters, or to delve into characterising the enzyme mechanism, in order to accurately address the role of the enzyme’s structure in the emergent epistasis.

## Methods

### Simple kinetic model simulations

We defined starting free energies (*G*) for each state as follows: E = 0.0 kcal mol^-1^, (E + S)^‡^ = 10.0 kcal mol^-1^, ES = −5.0 kcal mol^-1^, (ES)^‡^ = 11.0 kcal mol^-1^. We calculated microscopic rate constants for the wt using the following equations:

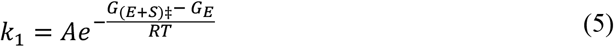

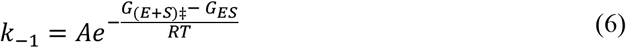

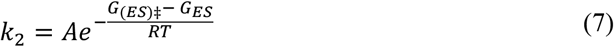

Where the pre-exponential factor *A* was approximated using transition state theory:

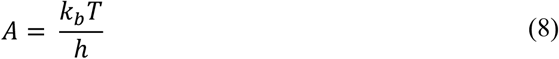

Where *k*_b_ is the Boltzmann constant and *h* is Plank’s constant. We note that this approximation permits *k*_1_ to approach 6.22 × 10^12^ M s^-1^, which exceeds the rate limit of diffusion by several orders of magnitude. However, for simplicity, we did not modify the constant for substrate binding calculation, and we do not expect results to differ greatly with a more accurate calculation of *A*.

We then computed kinetic parameters as follows:

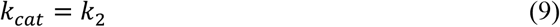

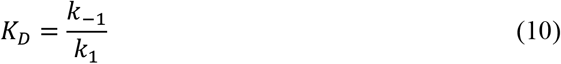

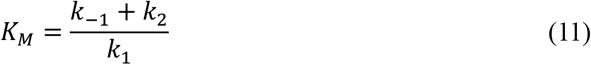

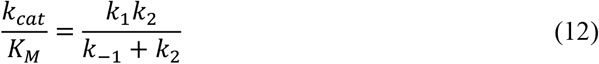

We generated 1,000 mutants *G* values for each parameter by iteratively drawing samples from a uniform distribution bounded between –2 and 2 and summing the wt *G* with the drawn sample.

Finally, we created 10^6^ “double mutants” by combinatorially combining all mutational effects from every single mutant. For the predicted kinetic parameter of the double mutant, we obtained the product of the wt kinetic parameter, the fold-change of mutant 1, and the fold-change of mutant 2. For the observed or “true” kinetic parameter, we first computed the microscopic rate constant of the double mutant by calculating the sum of free energy changes to each state in the reaction coordinate based on the single mutation effects. We used these rate constants to compute the observed kinetic parameter of the double mutant.

### Computation of mutational effects on epistasis in K_**M**_

In order to compute epistasis in *K*_M_ we used the following equation (derived in **S2 File**):

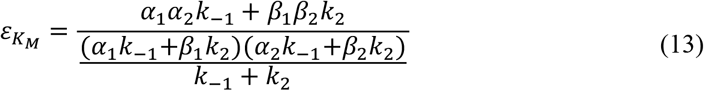

We then computed epistasis for combinations of *α*_1_, *α*_2_, *β*_1_, and *β*_2_ ranging from 0.01 to 100. For the initial computation *k*_-1_ = 62.1 s^-1^ and *k*_2_ = 11.5 s^-1^. Subsequent tests with variations in *k*_-1_ were performed at values relative to *k*_2_.

### Complex kinetic model simulations

We followed the steps as the more complex model, with additional energy states of (E + P)^‡^ = 9.0 kcal mol^-1^ and EP = −9.0 kcal mol^-1^ as well as the same definitions for *k*_1_, *k*_-1_, and *k*_2_, in addition to the new rate constants:

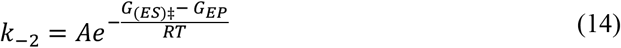

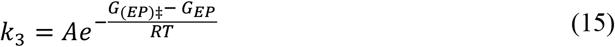

And updated kinetic parameters:

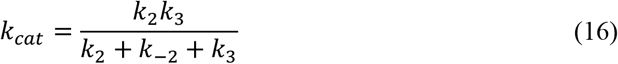

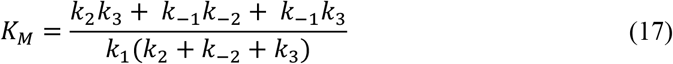

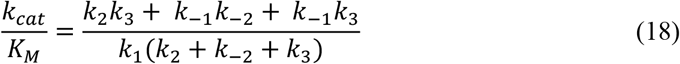

Where *K*_D_ remains the same as eq. 10

### Epistasis calculations in B. cerus β-lactamase I

Using equations 4.1, 4.2, and 4.3 we were able to compute *k*_cat_, *K*_M_, and catalytic efficiency using the microscopic rate constants reported in Gibson and Waley (1990) [38]. We found that computed *k*_cat_ of the single and double mutants was within 1.5-fold error of the measured *k*_cat_, however the computed *K*_M_ for K73R and the double mutant, as well as the computed catalytic efficiency for all mutants, was significantly different (>1.5-fold) than the measured parameters. We opted to use the computed single mutant kinetic parameters outlined in **Table 3**.

Null model fold-changes were calculated using the fold-changes for K73R and E166D which were computed by taking the ratio of the computed mutant parameter and the computed wt parameter. In contrast, corrected null model fold-changes were computed by first calculating the fold-change of each single mutant microscopic rate constant, then computing the expected microscopic rate constant for the double mutant by multiplying the wt constant with the two fold-changes. These expected values were also used for computation of specific epistasis for each microscopic rate constant. Next, we used equations 4.1–4.3 to compute expected kinetic parameters using the expected, not the observed, microscopic rate constants. The difference in these computed kinetic parameters relative to those observed and those computed using the null model provide the different values for epistasis.

## Software and data availability

We implemented the epistasis simulations using R (v4.5.0) supplemented with the libraries tidyverse (v2.0.0) and progress (v1.2.3). All simulations and software are available in the supporting information (**S1 and S3 Files**) as well as on GitHub (https://github.com/karolbuda/kinetic-epistasis-simulations). All mutant data are provided in a public repository [37].

## Acknowledgements

K.B. and N.T. are supported by the Natural Sciences and Engineering Research Council of Canada (NSERC)/Discovery Grants Program (RGPIN-2023-05135) and the Human Frontier Science Program (HFSP) Research Grant (RGP0054/2020). We thank Christopher Frøhlich and Adrian H. Bunzel, as well as all members of the Tokuriki lab, for discussions.

## Author contributions

Conceptualization: K.B. and N.T., Data curation: K.B., Formal analysis: K.B., Funding acquisition: N.T., Investigation: K.B. and N.T., Methodology: K.B., Project administration: N.T., Resources: K.B., Software: K.B., Supervision: N.T., Validation: K.B., Visualization: K.B., Writing – original draft: K.B., Writing – review and editing: K.B. and N.T.

## References

1. Card KJ, LaBar T, Gomez JB, Lenski RE. Historical contingency in the evolution of antibiotic resistance after decades of relaxed selection. Plos Biol. 2019;17: e3000397. doi:10.1371/journal.pbio.3000397

2. Xie VC, Pu J, Metzger BP, Thornton JW, Dickinson BC. Contingency and chance erase necessity in the experimental evolution of ancestral proteins. eLife. 2021;10: e67336. doi:10.7554/elife.67336

3. Starr TN, Picton LK, Thornton JW. Alternate evolutionary histories in the sequence space of an ancient protein. Nature. 2017;549: 409–413. doi:10.1038/nature23902

4. Bridgham JT, Ortlund EA, Thornton JW. An epistatic ratchet constrains the direction of glucocorticoid receptor evolution. Nature. 2009;461: 515–519. doi:10.1038/nature08249

5. Shah P, McCandlish DM, Plotkin JB. Contingency and entrenchment in protein evolution under purifying selection. Proc Natl Acad Sci. 2015;112: E3226–E3235. doi:10.1073/pnas.1412933112

6. Starr TN, Flynn JM, Mishra P, Bolon DNA, Thornton JW. Pervasive contingency and entrenchment in a billion years of Hsp90 evolution. Proc Natl Acad Sci United States Am. 2018;115: 4453–4458. doi:10.1073/pnas.1718133115

7. Natarajan C, Hoffmann FG, Weber RE, Fago A, Witt CC, Storz JF. Predictable convergence in hemoglobin function has unpredictable molecular underpinnings. Science. 2016;354: 336–339. doi:10.1126/science.aaf9070

8. Harms MJ, Thornton JW. Historical contingency and its biophysical basis in glucocorticoid receptor evolution. Nature. 2014;512: 203–207. doi:10.1038/nature13410

9. Kondrashov AS, Sunyaev S, Kondrashov FA. Dobzhansky–Muller incompatibilities in protein evolution. Proc National Acad Sci. 2002;99: 14878–14883. doi:10.1073/pnas.232565499

10. Jones AG, Bürger R, Arnold SJ. Epistasis and natural selection shape the mutational architecture of complex traits. Nat Commun. 2014;5: 3709. doi:10.1038/ncomms4709

11. Buda K, Miton CM, Tokuriki N. Pervasive epistasis exposes intramolecular networks in adaptive enzyme evolution. Nat Commun. 2023;14: 8508. doi:10.1038/s41467-023-44333-5

12. Johnson MS, Reddy G, Desai MM. Epistasis and evolution: recent advances and an outlook for prediction. BMC Biol. 2023;21: 120. doi:10.1186/s12915-023-01585-3

13. Chen JZ, Bisardi M, Lee D, Cotogno S, Zamponi F, Weigt M, et al. Understanding epistatic networks in the B1 β-lactamases through coevolutionary statistical modeling and deep mutational scanning. Nat Commun. 2024;15: 8441. doi:10.1038/s41467-024-52614-w

14. Poelwijk FJ, Tănase-Nicola S, Kiviet DJ, Tans SJ. Reciprocal sign epistasis is a necessary condition for multi-peaked fitness landscapes. J Theor Biol. 2011;272: 141–144. doi:10.1016/j.jtbi.2010.12.015

15. Visser JAGM de, Krug J. Empirical fitness landscapes and the predictability of evolution. Nat Rev Genet. 2014;15: 480–490. doi:10.1038/nrg3744

16. Starr TN, Thornton JW. Epistasis in protein evolution. Protein Sci. 2016;25: 1204–1218. doi:10.1002/pro.2897

17. Miton CM, Buda K, Tokuriki N. Epistasis and intramolecular networks in protein evolution. Curr Opin Struc Biol. 2021;69: 160–168. doi:10.1016/j.sbi.2021.04.007

18. Ramanathan A, Agarwal PK. Evolutionarily Conserved Linkage between Enzyme Fold, Flexibility, and Catalysis. PLoS Biol. 2011;9: e1001193. doi:10.1371/journal.pbio.1001193

19. Judge A, Sankaran B, Hu L, Palaniappan M, Birgy A, Prasad BVV, et al. Network of epistatic interactions in an enzyme active site revealed by large-scale deep mutational scanning. Proc Natl Acad Sci. 2024;121. doi:10.1073/pnas.2313513121

20. O’Rourke KF, Gorman SD, Boehr DD. Biophysical and computational methods to analyze amino acid interaction networks in proteins. Comput Struct Biotechnol J. 2016;14: 245–251. doi:10.1016/j.csbj.2016.06.002

21. Kurzbach D. Network representation of protein interactions: Theory of graph description and analysis. Protein Sci. 2016;25: 1617–1627. doi:10.1002/pro.2963

22. Kryazhimskiy S, Rice DP, Jerison ER, Desai MM. Global epistasis makes adaptation predictable despite sequence-level stochasticity. Science. 2014;344: 1519–1522. doi:10.1126/science.1250939

23. Kaltenbach M, Tokuriki N. Dynamics and constraints of enzyme evolution. J Exp Zoology Part B Mol Dev Evol. 2014;322: 468–487. doi:10.1002/jez.b.22562

24. Bershtein S, Segal M, Bekerman R, Tokuriki N, Tawfik DS. Robustness–epistasis link shapes the fitness landscape of a randomly drifting protein. Nature. 2006;444: 929–932. doi:10.1038/nature05385

25. Bloom JD, Arnold FH, Wilke CO. Breaking proteins with mutations: threads and thresholds in evolution. Mol Syst Biol. 2007;3: MSB4100119. doi:10.1038/msb4100119

26. Chen JZ, Fowler DM, Tokuriki N. Environmental selection and epistasis in an empirical phenotype–environment–fitness landscape. Nat Ecol Evol. 2022;6: 427–438. doi:10.1038/s41559-022-01675-5

27. Sarkisyan KS, Bolotin DA, Meer MV, Usmanova DR, Mishin AS, Sharonov GV, et al. Local fitness landscape of the green fluorescent protein. Nature. 2016;533: 397–401. doi:10.1038/nature17995

28. Otwinowski J, McCandlish DM, Plotkin JB. Inferring the shape of global epistasis. Proc National Acad Sci. 2018;115: 201804015. doi:10.1073/pnas.1804015115

29. Faure AJ, Martí-Aranda A, Hidalgo-Carcedo C, Beltran A, Schmiedel JM, Lehner B. The genetic architecture of protein stability. Nature. 2024;634: 995–1003. doi:10.1038/s41586-024-07966-0

30. Bendel AM, Faure AJ, Klein D, Shimada K, Lyautey R, Schiffelholz N, et al. The genetic architecture of protein interaction affinity and specificity. Nat Commun. 2024;15: 8868. doi:10.1038/s41467-024-53195-4

31. Sailer ZR, Harms MJ. Detecting High-Order Epistasis in Nonlinear Genotype-Phenotype Maps. Genetics. 2017;205: 1079–1088. doi:10.1534/genetics.116.195214

32. Park Y, Metzger BPH, Thornton JW. The simplicity of protein sequence-function relationships. Nat Commun. 2024;15: 7953. doi:10.1038/s41467-024-51895-5

33. Morrison AJ, Wonderlick DR, Harms MJ. Ensemble epistasis: thermodynamic origins of nonadditivity between mutations. Genetics. 2021;219: iyab105. doi:10.1093/genetics/iyab105

34. Michaelis L, Menten ML, Johnson KA, Goody RS. The Original Michaelis Constant: Translation of the 1913 Michaelis–Menten Paper. Biochemistry. 2011;50: 8264–8269. doi:10.1021/bi201284u

35. Bar-Even A, Noor E, Savir Y, Liebermeister W, Davidi D, Tawfik DS, et al. The Moderately Efficient Enzyme: Evolutionary and Physicochemical Trends Shaping Enzyme Parameters. Biochemistry. 2011;50: 4402–4410. doi:10.1021/bi2002289

36. Copley SD, Newton MS, Widney KA. How to Recruit a Promiscuous Enzyme to Serve a New Function. Biochemistry-us. 2022. doi:10.1021/acs.biochem.2c00249

37. Buda K, Tokuriki N. Supporting information for “Free energy perturbations in enzyme kinetic models reveal cryptic epistasis.” Zenodo. 2025. doi:10.5281/zenodo.17041247

38. Gibson RM, Christensen H, Waley SG. Site-directed mutagenesis of β -lactamase I. Single and double mutants of Glu-166 and Lys-73. Biochem J. 1990;272: 613–619. doi:10.1042/bj2720613

39. Rendón JL, Pardo JP. Time-Dependent Kinetic Complexities in Enzyme Assays: A Review. Biomolecules. 2025;15: 641. doi:10.3390/biom15050641

40. Fröhlich C, Bunzel HA, Buda K, Mulholland AJ, Kamp MW van der, Johnsen PJ, et al. Epistasis arises from shifting the rate-limiting step during enzyme evolution of a β-lactamase. Nat Catal. 2024; 1–11. doi:10.1038/s41929-024-01117-4

41. Acevedo-Rocha CG, Li A, D’Amore L, Hoebenreich S, Sanchis J, Lubrano P, et al. Pervasive cooperative mutational effects on multiple catalytic enzyme traits emerge via long-range conformational dynamics. Nat Commun. 2021;12: 1621. doi:10.1038/s41467-021-21833-w

42. Yu H, Dalby PA. Coupled molecular dynamics mediate long- and short-range epistasis between mutations that affect stability and aggregation kinetics. Proc Natl Acad Sci. 2018;115: E11043–E11052. doi:10.1073/pnas.1810324115

43. Markin CJ, Mokhtari DA, Sunden F, Appel MJ, Akiva E, Longwell SA, et al. Revealing enzyme functional architecture via high-throughput microfluidic enzyme kinetics. Science. 2021;373. doi:10.1126/science.abf8761

44. Markin CJ, Mokhtari DA, D. S, Doukov T, Sunden F, Cook JA, et al. Decoupling of catalysis and transition state analog binding from mutations throughout a phosphatase revealed by high-throughput enzymology. Proc Natl Acad Sci. 2023;120: e2219074120. doi:10.1073/pnas.2219074120

45. Muir DF, Asper GPR, Notin P, Posner JA, Marks DS, Keiser MJ, et al. Evolutionary-scale enzymology enables exploration of a rugged catalytic landscape. Science. 2025;388: eadu1058. doi:10.1126/science.adu1058

